# Polyphosphate synthesis in mast cell granules relies on V-ATPase activity supported by IP6K1

**DOI:** 10.64898/2026.08.07.742468

**Authors:** Manisha Mallick, Rashna Bhandari

## Abstract

Polyphosphate (polyP), a linear polymer of orthophosphate residues, is enriched in secretory granules in specialised mammalian cell types including platelets and mast cells. Although polyP released during activation and degranulation of these cells has been shown to promote blood clotting and inflammation, little is known about the mechanisms governing polyP synthesis in these granules. In mice, the loss of IP6K1, an enzyme that catalyses the production of 5-InsP_7_, has been shown to result in depletion of platelet polyP and impaired hemostasis. Here, we use the rat mast cell line RBL-2H3 as a model to study the regulation of polyP synthesis in secretory granules. By monitoring real-time polyP synthesis in isolated mast cell granules, we demonstrate that ATP is the substrate fuelling granule polyP production. By the use of inhibitors, we show that accumulation of polyP in granules requires an intact transmembrane proton gradient maintained by vacuolar H^+^ATPase (V-ATPase). In RBL-2H3 cells, depletion of IP6K1 led to a substantial reduction in cellular polyP levels and defective accumulation of polyP, serotonin, and tryptase inside granules. Cells with reduced IP6K1 showed a profound loss of granule acidification, correlating with downregulated levels of V_1_ subunits of V-ATPase. Adding back active or catalytically inactive IP6K1 rescued the expression of V-ATPase V_1_ subunits, reversed granule deacidification, and restored polyP levels in IP6K1-depleted cells. Together, these data unveil a role for IP6K1 in maintaining granule pH and thereby supporting polyP synthesis in mammals.

## INTRODUCTION

Polyphosphate (polyP), a linear biopolymer composed of three to several hundred orthophosphate residues linked by high-energy phosphoanhydride bonds, supports vital functions across various forms of life [1–3]. In prokaryotes and unicellular eukaryotes, polyP contributes to phosphate homeostasis, stress response, and to virulence in the case of pathogenic microorganisms [1, 4–6]. Bacteria synthesize long chains of polyP from ATP via the enzyme polyphosphate kinase 1 (PPK1), whereas in most unicellular eukaryotes, production of medium chain length polyP (∼60-100 Pi units) is carried out by the vacuolar transport chaperone (VTC) complex located on the membranes of vacuoles and acidocalcisomes [7–10]. Metazoans, which do not possess homologs of PPK1 or VTC, synthesize low levels of polyP by an enzyme that is yet to be definitively identified. In *Drosophila,* larval development and hemolymph clotting are regulated by polyP [11]. In mammalian cells, polyP is present in the nucleus, cytoplasm, and mitochondria, and is especially enriched in lysosome related organelles (LROs) that resemble the acidocalcisomes found in protists [12–17]. In mammalian platelets and mast cells, polyP stored in secretory granules is released upon degranulation and acts as a procoagulant and proinflammatory mediator [13, 15, 18–21]. In astrocytes and neurons, polyP is stored in secretory vesicles, which upon release, regulates neuronal excitability [22–25]. Additionally, in mammals, polyP also participates in bone mineralisation, cell differentiation and proliferation, stress response, energy metabolism, and formation of biomolecular condensates [26–38].

In the budding yeast *S. cerevisiae*, vacuolar polyP synthesis is allosterically upregulated by binding of the inositol pyrophosphate 5-diphosphoinositol pentakisphosphate (5-InsP_7_) to the SPX (Syg1/Pho81/Xpr1) domain of VTC4, the catalytic subunit of the VTC complex [39–41]. Binding of 5-InsP_7_ to the VTC4 SPX domain causes a conformational change in the VTC complex that allows the coupled synthesis and translocation of nascent polyP into the vacuole lumen [41–43]. Although mammals do not possess a VTC4 homolog, inositol hexakisphosphate kinase (IP6K1), an enzyme that synthesises 5-InsP_7,_ has been shown to support polyP synthesis in mitochondria and platelet dense granules [44, 45]. In isolated mitochondria, polyP synthesis requires an intact transmembrane proton gradient and depends on the activity of F_0_F_1_ ATP synthase, which may serve directly as a polyP synthase [45–47]. Mitochondria isolated from cells and mice deficient in IP6K1 show a marked reduction in polyP synthesis, which can be rescued by expression of active IP6K1 capable of synthesizing 5-InsP_7_, but not by catalytically inactive IP6K1 [45]. Additionally, IP6K1 was shown to support mitochondrial respiration independently of its catalytic activity. We have previously reported that *Ip6k1* knockout mice display a substantial reduction in platelet polyP levels, which is associated with impaired hemostasis in these mice [44]. The mechanism by which IP6K1 supports polyP synthesis in platelet granules, and whether this is dependent on 5-InsP_7_ synthesis, is still unclear.

In this study, we have used the rat basophilic leukaemia cell line RBL-2H3, which resemble mast cells [15], as a tractable model to explore polyP synthesis in LROs and investigate the relationship between IP6K1 and granule polyP. Our data show that polyP is synthesised in isolated RBL-2H3 granules by utilising ATP as a phosphate source. An intact proton gradient across the granule membrane, maintained by activity of the vacuolar-ATPase (V-ATPase), is indispensable for polyP production in granules. We further demonstrate that IP6K1 depletion in RBL-2H3 cells reduces granule acidification by downregulating the expression of V_1_ subunits of V-ATPase. When catalytically active or inactive IP6K1 are reintroduced into *Ip6k1* knockdown RBL-2H3 cells, they can restore V-ATPase expression, granule acidification, and polyP levels. Our data provides the first definitive evidence of polyP synthesis in isolated mammalian LROs, and offers a mechanistic explanation for the regulation of granule polyP by IP6K1.

## RESULTS

### PolyP is synthesised in isolated RBL-2H3 granules

To study polyP in mast cells we used the rat basophilic leukaemia cell line RBL-2H3, which possesses mast cell-like properties including cell surface IgE receptors and cytoplasmic granules containing inflammatory mediators such as bioamines and hydrolases [48]. RBL-2H3 cells have been shown to store polyP in cytoplasmic granules that display homology with acidocalcisomes [15]. Using an in situ protein overlay assay, we first confirmed the presence of polyP-containing granules in RBL-2H3 cells. We generated a construct expressing GST-tagged polyP binding domain (PPBD) of *E. coli* exopolyphosphatase, which has been shown to specifically bind polyP [15, 49–51]. GST-PPBD was added to fixed and permeabilised RBL-2H3 cells, followed by detection of GST using immunofluorescence. We observed distinct cytoplasmic granular staining in RBL-2H3 cells probed with GST-PPBD, whereas GST, used as a control, showed diffuse staining throughout the cell (Fig. 1A). Protein mediators such as tryptase and β-hexosaminidase are transported from the ER via the Golgi to mast cell granules, whereas biogenic amines including serotonin and histamine are synthesized in the cytoplasm and transported into granules [52, 53]. Co-staining RBL-2H3 cells with GST-PPBD and an antibody to detect serotonin, confirmed that polyP co-localises with serotonin-containing granules as reported previously [15] (Fig. 1B).

**Figure 1.**
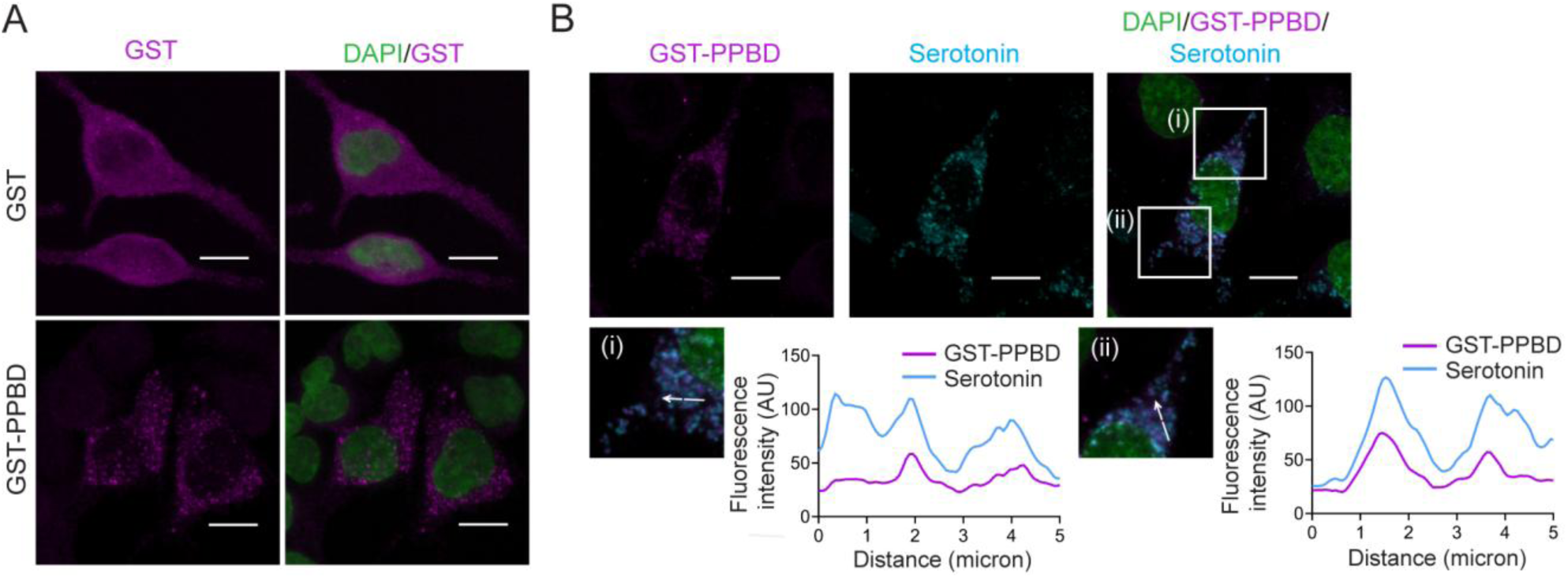
Localisation of polyP in RBL-2H3 cells. **(A)** In situ protein overlay assay to detect polyP in RBL-2H3 cells. Representative images show fixed and permeabilised RBL-2H3 cells incubated with GST-PPBD (which binds polyP) or GST (control), followed by detection with GST antibody (magenta); DAPI (green) marks the nucleus. n = 24 cells for GST and 20 cells for GST-PPBD over two independent experiments. **(B)** Immunofluorescence analysis of RBL-2H3 cells following in situ protein overlay, shows GST-PPBD to detect polyP (magenta), serotonin (cyan), and DAPI to mark nuclei (green). Confocal microscopy images in (A and B), showing individual channels and an overlay, were captured with a Leica TCS SP8 confocal microscope using a 63x, 1.4 NA oil-immersion objective, and are presented as z-stacks with xy dimensions as a maximum intensity projection (MIP); scale bar: 10 μm. White boxes (i) and (ii) in (B) mark the regions in the overlay image that are zoomed in and presented below. Line graphs show the fluorescence intensity for serotonin (cyan) and GST-PPBD (magenta) measured along the indicated white arrows. Pearson’s correlation coefficient to measure co-localisation of polyP and serotonin in (B) is 0.66 ± 0.15 (n = 18 cells).

To investigate whether polyP is synthesized within granules or, like biogenic amines, is imported into granules from the cytoplasm, we conducted polyP synthesis assays in organelle fractions isolated from RBL-2H3 cells. We used differential centrifugation to isolate membrane-intact organelles including mitochondria, the endomembrane system (ER and Golgi), and granules, and verified organelle enrichment via western blotting using specific markers (Fig. S1A, B) [54, 55]. VAMP8, the v-SNARE that is known to mark serotonin-containing granules [56], was enriched in the granule fraction (Fig. S1A, B). These fractions were subjected to *ex vivo* polyP synthesis by measuring fluorescence of polyP-bound DAPI in real time, a method used previously to detect polyP synthesis in isolated mitochondria [45]. We incubated the mitochondria, endomembrane system, and granule fractions with ATP (1 mM) and inorganic phosphate (Pi; 5 mM) in the presence of DAPI, and continuously monitored the change in DAPI fluorescence (ΔF), corresponding to polyP accumulation over time. As a control, we confirmed that ATP and Pi alone do not contribute to DAPI fluorescence (Fig. S1C). The organelle fractions did not show any substantial change in fluorescence over time in the absence of ATP and Pi (Fig. 2A), whereas in the presence of these phosphate sources, the granule fraction showed a significantly higher ΔF compared with the mitochondria and the endomembrane system (Fig. 2B, C). These data clearly show that isolated granules from RBL-2H3 cells are capable of polyP synthesis.

**Figure 2.**
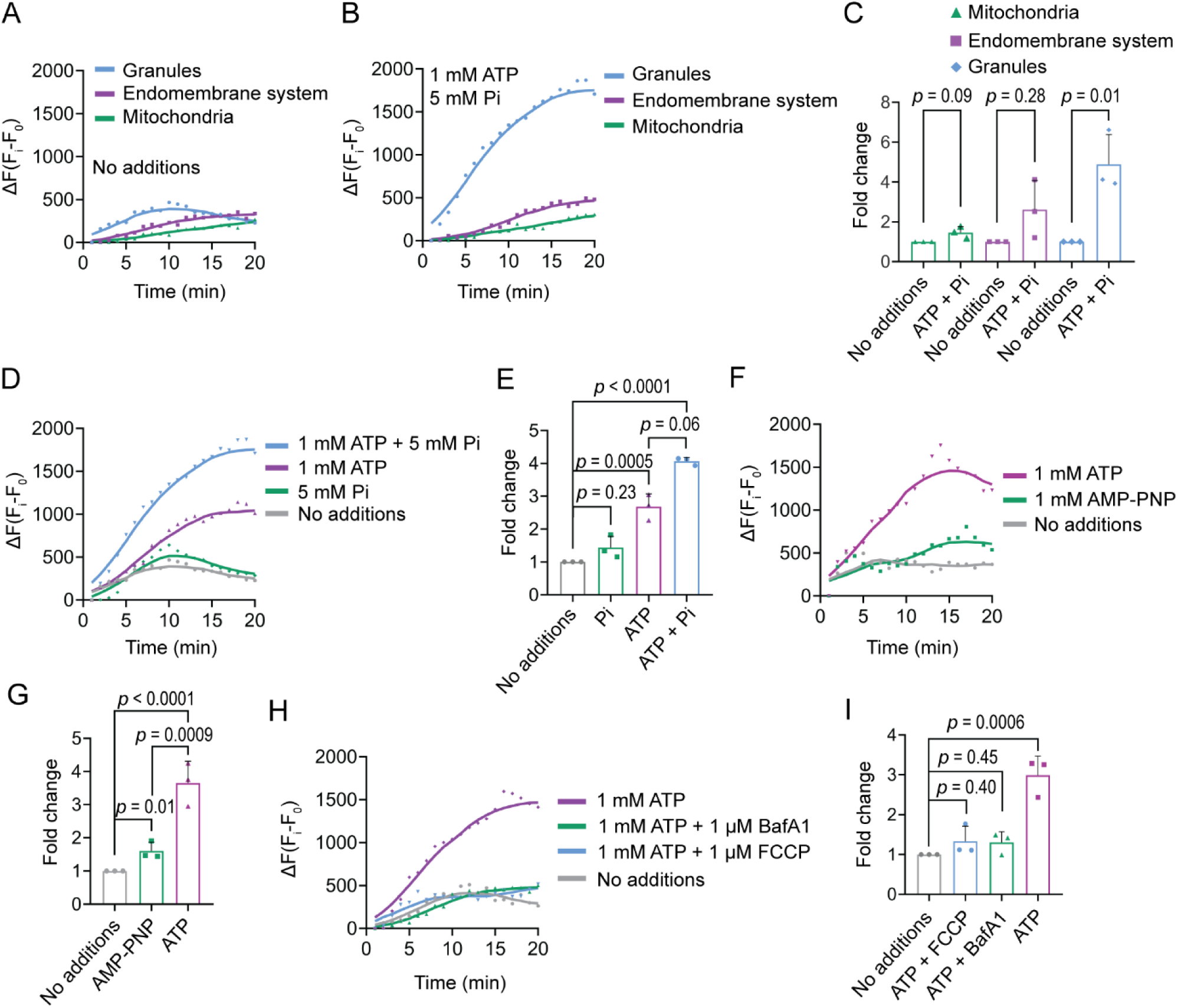
PolyP synthesis in granules isolated from RBL-2H3 cells. **(A, B)** Line graphs represent the change in DAPI fluorescence (ΔF) over time as a measure of polyP synthesis in isolated mitochondria, endomembrane system, and granules, in the absence (A) or in the presence (B) of Pi (5 mM) and ATP (1 mM). **(C)** Quantification of (A, B). Bar graphs represent the fold change in polyP accumulated over 20 min in isolated mitochondria, endomembrane system, and granules, in the presence of ATP (1 mM) and Pi (5 mM) (shown in B), over their respective controls (shown in A). **(D)** Line graphs represent ΔF over time in isolated granules in the absence or presence of ATP (1 mM), or Pi (5 mM), or both. The same representative line graph is used in (B) and (D) for polyP synthesis in the isolated granule fraction in the presence of ATP (1 mM) and Pi (5 mM). **(E)** Quantification of (D). Bar graphs represent the fold change in polyP levels in isolated granules in the presence of Pi, ATP, or both, over the control. **(F)** Line graphs represent ΔF over time in isolated granules in the absence or presence of ATP (1 mM) or AMP-PNP (1 mM). **(G)** Quantification of (F). Bar graphs represent the fold change in polyP levels in isolated granules in the presence of ATP or AMP-PNP, over the control. **(H)** Line graphs represent ΔF over time in isolated granules in the absence or presence of ATP (1 mM), with or without the addition of BafA1 (1 µM), or FCCP (1 µM). **(I)** Quantification of (H). Bar graphs represent the fold change in polyP levels in isolated granules in the presence of ATP (1 mM), with or without the addition of BafA1 or FCCP, over the control. Line graphs in (A, B, D, F, and H) were generated in GraphPad Prism 8 using the curve smoothening function, averaging 4 values on each side, and using a 0th-order smoothing polynomial. Area under the curve (AUC) for each line graph of the test samples was normalised to the control (no additions), and the fold change (mean ± S.D.) is represented in the bar graphs in (C, E, G, and I). To determine *p* values, the fold change values were transformed to log_2_ and subjected to a one-sample *t*-test against a theoretical mean of 0 (C), or one-way ANOVA followed by Tukey’s multiple comparisons test (E, G, and I); N = 3.

We have previously shown that isolated mitochondria can synthesize polyP from Pi in the presence of substrates of the electron transport chain (ETC), with ATP enhancing Pi-driven mitochondrial polyP synthesis [45]. In the absence of ETC substrates, we observed no significant mitochondrial polyP synthesis in the presence of Pi and ATP (Fig. 2B, C). To identify the primary substrate for polyP synthesis in granules, we tracked *ex vivo* polyP synthesis in isolated granules in the presence of either ATP or Pi, and found that ATP alone was sufficient to initiate polyP synthesis (Fig. 2D, E). Although Pi alone failed to trigger *de novo* synthesis, the presence of Pi amplified ATP-dependent polyP accumulation (Fig. 2D, E), suggesting that Pi may have a secondary role in polyP production. AMP-PNP, a non-hydrolysable analogue of ATP, did not support granule polyP synthesis to the same extent as ATP (Fig. 2F, G), reinforcing our conclusion that ATP is the source of phosphate for polyP synthesis in isolated granules. In isolated yeast vacuoles, VTC-dependent polyP synthesis and translocation into the lumen is dependent on the electrochemical proton gradient across the vacuole membrane, which is established and maintained by the vacuolar H^+^-ATPase (V-ATPase) [57]. As mast cell granules also maintain a V-ATPase-dependent proton gradient across their membrane, with the granule lumen being acidic compared with the cytoplasm [58, 59], we examined the importance of granule acidification in polyP synthesis. Bafilomycin A1 (BafA1), an inhibitor of V-ATPase activity, and FCCP (carbonyl cyanide-ptrifluoromethoxyphenylhydrazone), a proton uncoupler, are known to disrupt the proton gradient across the granule membrane [60, 61]. In isolated granules, the addition of BafA1 (1 µM) or FCCP (1 µM) abrogated ATP-driven polyP production (Fig. 2H, I). Together, these data reveal that granule polyP synthesis uses ATP as a substrate and relies on an intact electrochemical membrane potential. Interestingly, these features of polyP synthesis are conserved between mammalian mast cell granules and budding yeast vacuoles despite the lack of conservation of the polyP synthase.

### Depletion of IP6K1 reduces granule polyP levels

We have previously shown that in mice, the loss of IP6K1, an enzyme responsible for the synthesis of 5-InsP_7_ from InsP_6_, leads to a reduction in platelet polyP levels [44]. Consequently, these mice display impaired blood clotting and reduced formation of neutrophil-platelet aggregates, both processes known to be influenced by polyP released during platelet degranulation [44, 62]. Platelets, derived from megakaryocytes, share their lineage with mast cells – both cell types arise from a common myeloid progenitor [63, 64]. The regulation of granule polyP levels by IP6K1 may therefore be conserved between platelets and RBL-2H3 cells. To examine this possibility, we depleted IP6K1 in RBL-2H3 cells using shRNA targeted to rat *Ip6k1*. We obtained two lines (sh*Ip6k1*-1 and sh*Ip6k1*-2) in which the extent of knockdown of IP6K1 expression was 70-90% compared with the non-targeted control (shNT) RBL-2H3 line (Fig. 3A). We extracted polyP from these cell lines using acid phenol [65], and measured it using the polyP-DAPI fluorescence method we have described earlier [45]. PolyP levels were reduced by more than 80% in both *Ip6k1* knockdown lines compared with non-targeted control RBL-2H3 cells (Fig. 3B). Next, we co-stained RBL-2H3 cells to detect polyP and serotonin containing granules, and observed a significant loss in fluorescence signal intensity for both markers in IP6K1 depleted RBL-2H3 cells (Fig. 3C-E). These data confirm that the effect of IP6K1 depletion on granule polyP is preserved in platelets and mast cells.

**Figure 3.**
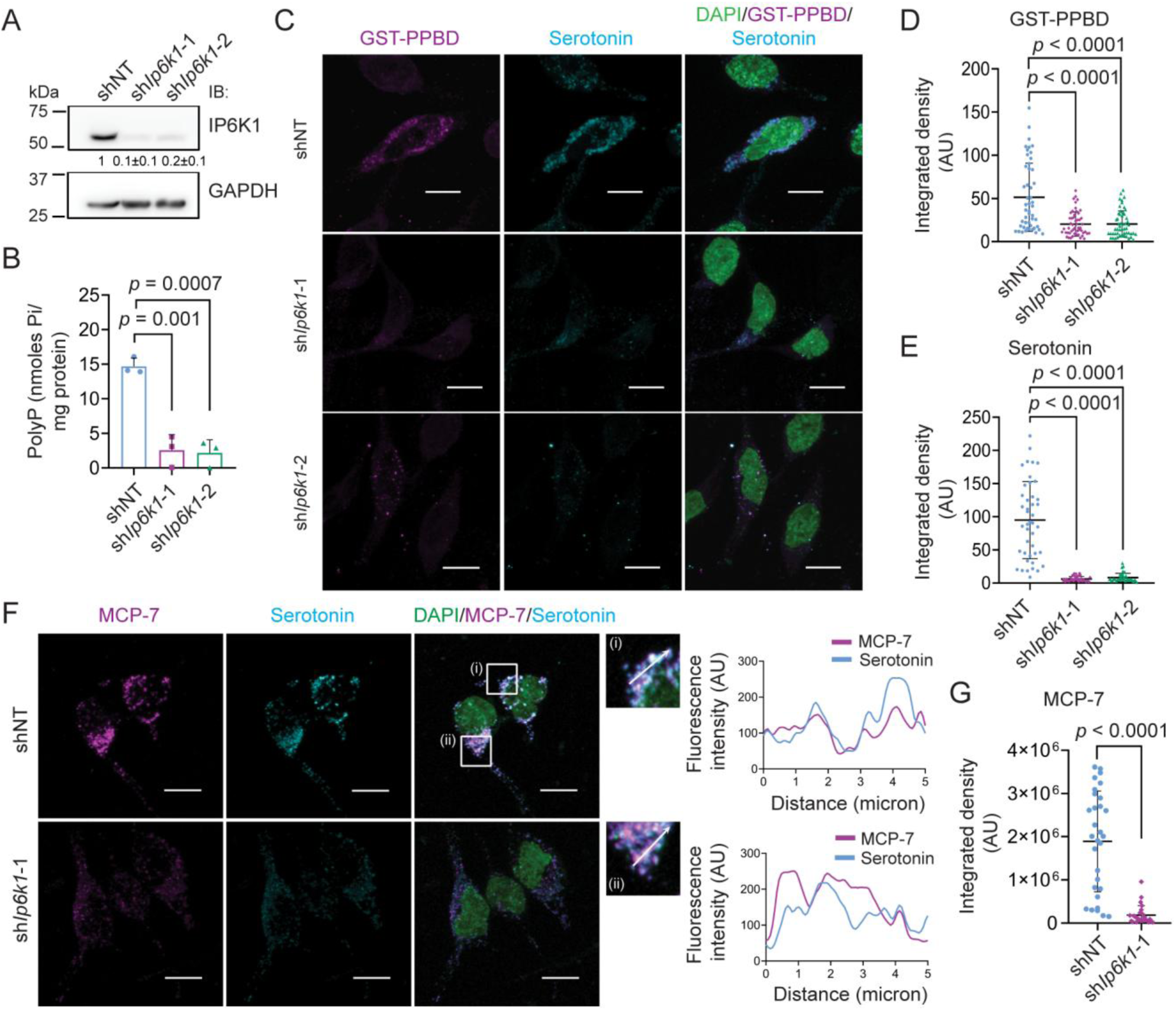
Depletion of IP6K1 affects polyP, serotonin and tryptase accumulation in RBL-2H3 cells. **(A)** Immunoblots show endogenous IP6K1 levels in non-targeted control (shNT) and IP6K1 knockdown (sh*Ip6k1*-1 and sh*Ip6k1*-2) RBL-2H3 cell lines. The extent of IP6K1 expression (mean ± S.D.) in knockdown lines compared with the non-targeted control is indicated (N = 3). **(B)** Bar graphs (mean ± S.D.) represent polyP levels in the indicated RBL-2H3 cell lines (N = 3). *p* values were determined using a two-tailed unpaired Student’s *t*-test. **(C)** Immunofluorescence analysis of indicated RBL-2H3 cell lines following in situ protein overlay, shows GST-PPBD to detect polyP (magenta), serotonin (cyan), and DAPI to mark nuclei (green). **(D, E)** Quantification of (C). Scatter dot plots (with mean ± S.D.) represent GST-PPBD (D) and serotonin (E) fluorescence signals as integrated density per cell (in arbitrary units, AU) in the indicated RBL-2H3 cell lines. Sample sizes were n = 50, 54 and 55 for GST-PPBD and n = 43, 43 and 41 for serotonin in shNT, sh*Ip6k1*-1, and sh*Ip6k1*-2 RBL-2H3 cell lines, respectively. Data were compiled from three independent experiments. The adjusted *p* values were estimated using a Kruskal-Wallis test with Dunn’s multiple comparisons test. **(F)** Immunofluorescence analysis in the indicated RBL-2H3 cell lines shows MCP-7 (tryptase; magenta), serotonin (cyan), and DAPI to mark nuclei (green). White boxes (i) and (ii) mark the regions in the overlay image that are zoomed in and shown on the right. Line graphs show the fluorescence intensity for serotonin (cyan) and MCP-7 (magenta) measured along the indicated white arrows. Pearson’s correlation coefficient to measure co-localisation of serotonin and MCP7 is 0.54 ± 0.16 (n = 20 cells). **(G)** Scatter dot plots (with mean ± S.D.) represent MCP-7 fluorescence signals as integrated density per cell (in arbitrary units, AU) in shNT (n = 31) and sh*Ip6k1*-1 (n = 30) RBL-2H3 cell lines. *p* value was determined using a two-tailed Mann-Whitney test. Data were compiled from three independent experiments. Confocal microscopy images in (C) and (F), showing individual channels and an overlay, were captured with a Leica TCS SP8 confocal microscope using a 63x, 1.4 NA oil-immersion objective, and are presented as z-stacks with xy dimensions as a maximum intensity projection (MIP); scale bar: 10 μm.

To determine whether the loss of IP6K1 affects only polyP and serotonin, or whether other granule mediators are also impacted, we stained RBL-2H3 cells to detect the protease tryptase (MCP-7), which is known to accumulate in mast cell secretory granules [53, 66]. We observed co-localisation of tryptase with serotonin containing granules in shNT cells, and noted a drastic reduction in tryptase staining in *Ip6k1* knockdown cells (Fig. 3F, G). Interestingly, the levels of tryptase were unchanged in cells depleted for IP6K1 (Fig. S2A, B), revealing that IP6K1 does not regulate tryptase synthesis but instead may influence its trafficking to, or retention within, granules. In a similar manner, IP6K1 has previously been shown to support the trafficking and accumulation of the digestive enzymes pepsinogen C (PGC) and gastric lipase F (LIPF) in secretory granules of gastric chief cells without affecting the expression levels of these enzymes [67]. We also examined whether IP6K1 depletion affects another mast cell granule mediator – the lysosomal hydrolase β-hexosaminidase [53, 66]. The cellular levels, sub-cellular distribution, and enzymatic activity of β-hexosaminidase was unaltered in sh*Ip6k1* RBL-2H3 cells (Fig. S2C-G). Together, these data suggest that IP6K1 may support the accumulation of polyP, serotonin and tryptase in mast cell granules via a common underlying regulatory mechanism.

### IP6K1 depletion impairs granule acidification

In concordance with our data on the impact of IP6K1 depletion on granule mediators in RBL-2H3 cells, a previous study has shown that treatment of mast cells with BafA1 to abrogate granule acidification reduced tryptase activity but did not affect the activity of β-hexosaminidase [68]. Additionally, the import of serotonin from the cytoplasm into granules by vesicular monoamine transporters (VMATs) is known to rely on an intact proton gradient generated by V-ATPase [52]. PolyP synthesis and retention in isolated RBL-2H3 granules also requires an intact protein gradient dependent on the activity of V-ATPase (Fig. 2H and I) [15]. Together, these data led us to hypothesize that IP6K1 may act to sustain mast cell granule acidification by modulating V-ATPase. We therefore examined granule pH in control and *Ip6k1* knockdown RBL-2H3 cells using LysoSensor Green DND-189. This acidotropic probe accumulates in lysosome-related organelles, and its fluorescence is inversely dependent on the pH in the range of 4.5-6.0. We observed a drastic reduction in the fluorescence intensity of LysoSensor in *Ip6k1* knockdown RBL-2H3 cells, indicative of compromised granule acidification in these cells (Fig. 4A, B). Next, we examined the levels of V-ATPase, a multimeric complex composed of a cytoplasmic V_1_ domain responsible for ATP hydrolysis, and a membrane-embedded V_0_ domain engaged in proton translocation [69, 70]. In mammals, the V_1_ subcomplex is made from eight different subunits (A through H) with a composition ratio of A_3_B_3_CDE_3_FG_3_H, and the V_0_ component contains six core subunits with the ratio ac_8-9_c”def and two additional subunits ATP6AP1 and ATP6AP2. Immunoblotting to detect V1A and V1B, which comprise the catalytic A_3_B_3_ hexamer in the V_1_ domain, revealed a substantial drop in the levels of these subunits in IP6K1-depleted RBL-2H3 cells (Fig. 4C, D, S3A, B). The levels of subunit V1D of the central stalk and subunit V1E of the peripheral stalk are also significantly lower in *Ip6k1* knockdown cells, suggesting that the entire V_1_ complex is downregulated in the absence of IP6K1 (Fig. 4C, D, S3C, D). Interestingly, there was no effect of IP6K1 depletion on the levels of the V_0_ subunit a (V0A1; Fig. 4E, S3E). The V_0_ subcomplex is assembled in the ER and trafficked through the Golgi to vesicles, whereas the V_1_ subcomplex is assembled in the cytoplasm and anchors to the V_0_ domain based on regulatory cues that control intra-vesicle pH [71]. The differential regulation of V_1_ and V_0_ domains by IP6K1 is consistent with their distinct spatial origins. As subunits of the V_1_ subcomplex are known to undergo proteasomal degradation [72, 73], we tested the effect of the proteasomal inhibitor MG132 on the levels of the V_1_ subunits in control and *Ip6k1* knockdown RBL-2H3 cells. Treatment with M132 reversed the decrease in V1A, VIB and V1E subunits in sh*Ip6k1* cells, whereas it had no significant impact on V1D levels (Fig S3F-J).

**Figure 4.**
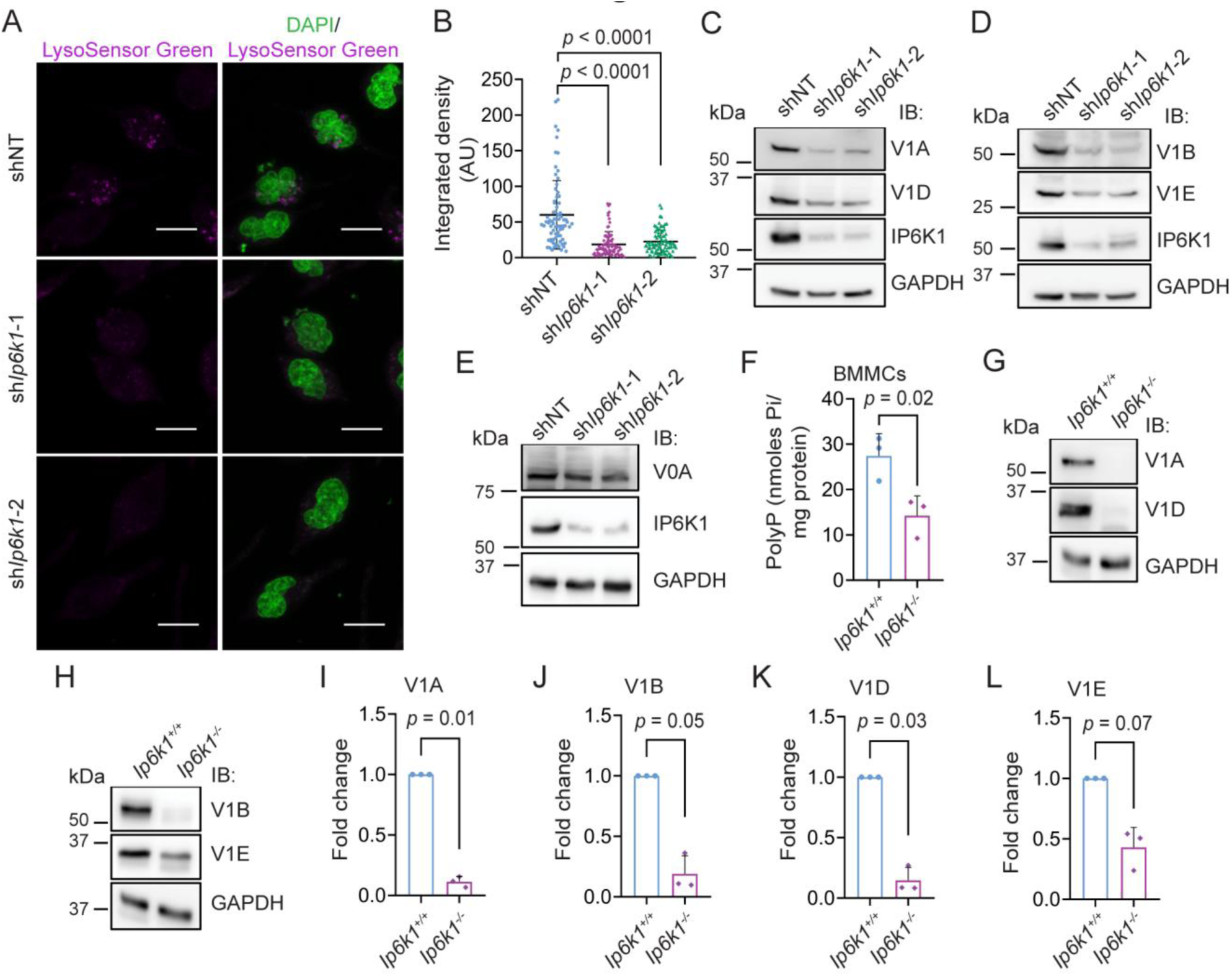
Loss of IP6K1 impairs granule acidification in RBL-2H3 cells. **(A)** Live cell fluorescence images of the indicated RBL-2H3 cell lines stained with LysoSensor Green (magenta) and DAPI to mark nuclei (green). Images showing individual channels and an overlay were captured with a Leica TCS SP8 confocal microscope using a 63x, 1.4 NA oil-immersion objective, and are presented as z-stacks with xy dimensions as a maximum intensity projection (MIP); scale bar: 10 μm. **(B)** Quantification of (A). Scatter dot plots (with mean ± S.D.) represent LysoSensor Green fluorescence signals as integrated density per cell (in arbitrary units, AU) in shNT (n = 81), sh*Ip6k1*-1 (n = 86), and sh*Ip6k1*-2 (n = 87) RBL-2H3 cells. Adjusted *p* values were determined using a Kruskal-Wallis test with Dunn’s multiple comparisons test. Data were compiled from three independent experiments. **(C-E)** Immunoblots show the levels of V-ATPase subunits V1A and V1D (C); V1B and V1E (D); and V0A (E) in the indicated RBL-2H3 cell lines. The blots also show levels of IP6K1 and GAPDH as a loading control. **(F)** Bar graphs (mean ± S.D.) represent polyP levels in bone marrow derived mast cells (BMMCs) from *Ip6k1^+/+^* and *Ip6k1^-/-^* mice (N = 3). The *p* value was determined using a two-tailed unpaired Student’s *t*-test. **(G, H)** Immunoblots show the levels of V-ATPase subunits V1A and V1D (G); V1B and V1E (H) in BMMCs from *Ip6k1^+/+^* or *Ip6k1^-/-^* mice. GAPDH was used as a loading control. **(I-L)** Quantification of (G, H). Bar graphs (mean ± S.D.) represent the levels of V1A (I), V1B (J), V1D (K) and V1E (L) in BMMCs from *Ip6k1^-/-^* mice compared with *Ip6k1^+/+^* mice (N = 3). The fold change values were transformed to log_2_, and the *p* values were determined using a one-sample *t*-test against a theoretical mean of 0.

To confirm that the regulation of polyP by IP6K1 is also conserved in primary mast cells, we extracted hematopoietic stem cells from the bone marrow of *Ip6k1^+/+^* and *Ip6k1^-/-^* mice and induced their differentiation into mast cells by culturing them in the presence of interleukin-3 (IL3; 10 µg/ml) for 21 days. Using flow cytometry to detect cell surface expression of the high-affinity IgE receptor (FcεRIα subunit), we confirmed that approximately 70% of the hematopoietic stem cells from both *Ip6k1^+/+^*and *Ip6k1^-/-^* mice had differentiated into mast cells (Fig. S3K). We observed a nearly 50% reduction in polyP levels in bone marrow-derived mast cells (BMMCs) from *Ip6k1^-/-^* mice compared with *Ip6k1^+/+^* mice (Fig. 4F). There was also a dramatic reduction in the levels of V_1_ subunits in BMMCs lacking IP6K1 (Fig. 4G-L), clearly recapitulating our observations in IP6K1-depleted RBL-2H3 cells. Collectively, these data reveal a role for IP6K1 in facilitating granule acidification by directly or indirectly preventing proteasomal degradation of the cytoplasmic subunits of V-ATPase in mast cells.

### IP6K1 supports granule acidification independent of its catalytic activity

So far we have seen that polyP synthesis in isolated mast cell granules requires an intact proton gradient across the granule membrane, and that the loss of IP6K1 in mast cells negatively impacts granule acidification by downregulating the levels of V-ATPase V_1_ subunits. Next, we examined whether IP6K1 supports granule acidification and polyP synthesis via generating 5-InsP_7_ or by a catalytic activity-independent mechanism. We reintroduced kinase-active or inactive (K226A/S334A) IP6K1 [74] in sh*Ip6k1*_1 RBL-2H3 cells (Fig. S4A) and compared the expression of V-ATPase V_1_ subunits in these cell lines alongside non-targeted control cells (Fig. 5A, B). Overexpression of either active or inactive IP6K1 rescued the expression levels of V-ATPase V_1_ subunits in IP6K1 depleted cells (Fig. S4B-E). As V-ATPase is the primary regulator of granule acidification, we imaged cells loaded with LysoSensor Green DND-189 to determine granule pH (Fig. 5C). Expression of either active or inactive IP6K1 resulted in partial recovery of granule acidification in *Ip6k1* knockdown RBL-2H3 cells, reflecting the restoration of V-ATPase levels (Fig. 5C, D). Finally, we examined whether IP6K1 overexpression can restore polyP levels in sh*Ip6k*1 RBL-2H3 cells. In keeping with the ability of IP6K1 to support V-ATPase expression and function independent of its catalytic activity, we observed that expression of active or inactive IP6K1 could restore polyP in IP6K1 depleted RBL-2H3 cells (Fig. 5E). To confirm that IP6K1 does indeed support cellular polyP levels independent of its ability to synthesize 5-InsP_7_, we treated RBL-2H3 cells with the pan IP6K inhibitor TNP [N2-(m-trifluorobenzyl), N6-(p-nitrobenzyl) purine], and confirmed the depletion of cellular InsP_7_ (Fig. S4F). TNP treatment did not lead to any change in total polyP levels in RBL-2H3 cells compared with the DMSO treated control (Fig. S4G). Together, our data suggest that IP6K1 acts independently of its catalytic activity to support the expression of V-ATPase V_1_ subunits, thereby ensuring granule acidification and polyP synthesis.

**Figure 5.**
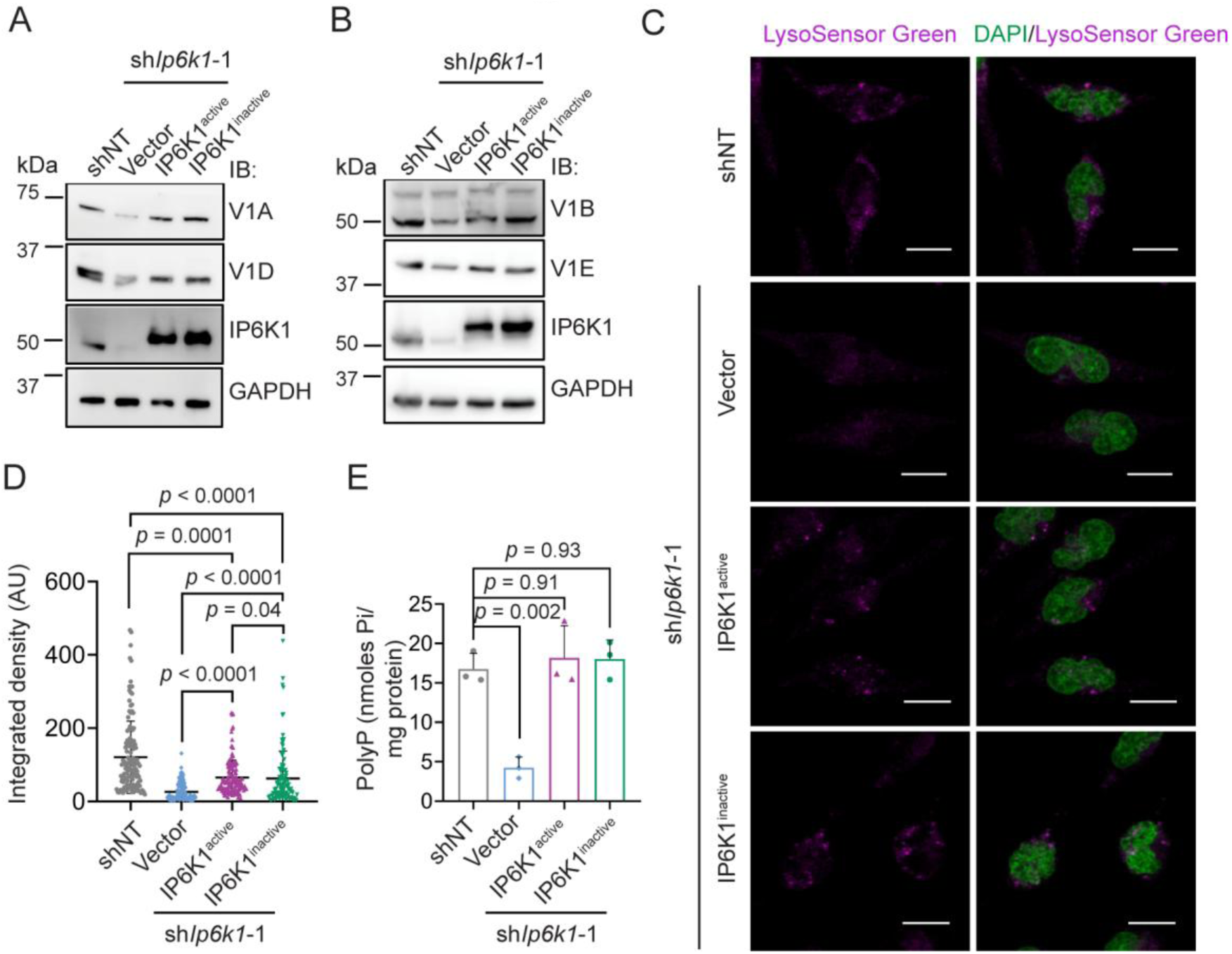
IP6K1 regulates granule acidification and polyP independent of its catalytic activity. **(A, B)** Immunoblots show the levels of V-ATPase subunits V1A and V1D (A); V1B and V1E (B) in non-targeted control (shNT) cells, sh*Ip6k1*-1 cells transfected with vector alone, or sh*Ip6k1*-1 RBL-2H3 cells stably expressing V5-tagged IP6K1 that is active (IP6K1^active^) or catalytically inactive (IP6K1^inactive^; K226A/S334A). The blots also show levels of IP6K1 and GAPDH as a loading control. **(C)** Live cell fluorescence images of the indicated RBL-2H3 cell lines stained with LysoSensor Green (magenta) and DAPI to mark nuclei (green). Images showing individual channels and an overlay were captured with a Leica TCS SP8 confocal microscope using a 63x, 1.4 NA oil-immersion objective, and are presented as z-stacks with xy dimensions as a maximum intensity projection (MIP); scale bar: 10 μm. **(D)** Quantification of (C). Scatter dot plots (with mean ± S.D.) represent LysoSensor Green fluorescence signals as integrated density (in arbitrary units, AU) per cell in shNT (n = 128), sh*Ip6k1*-1 with vector (n = 112), and sh*Ip6k1*-1 overexpressing IP6K1^active^ (n = 132) or IP6K1^inactive^ (n = 108) RBL-2H3 cells. Data were compiled from three independent experiments. *p* values were determined using a Kruskal-Wallis test with Dunn’s multiple comparisons test. **(E)** Bar graphs (mean ± S.D.) represent polyP levels in the indicated RBL-2H3 cell lines (N = 3). Adjusted *p* values were determined using one-way ANOVA followed by a Tukey’s multiple comparisons test.

## DISCUSSION

Although mammalian polyP is known to be enriched in platelet and mast cell granules, the mechanisms of polyP synthesis and regulation in these LROs remain poorly understood. Our study shows that isolated mast cell granules can synthesise polyP in an ATP-dependent manner. We observe that the electrochemical gradient across the granule membrane, and V-ATPase activity which maintains this gradient, are essential for polyP synthesis in isolated granules. Additionally, our work has unveiled a role for IP6K1 in polyP production by mast cell granules. The depletion of IP6K1 in mast cells impaired vesicular acidification, thereby causing loss of granule polyP. Interestingly, the accumulation of other granule mediators, such as serotonin and tryptase, was also disrupted in the absence of IP6K1. We see that IP6K1 acts independently of its catalytic activity to maintain levels of the V_1_ sub-complex of V-ATPase, thereby supporting granule acidification and the accumulation of granule mediators including polyP (Fig. 6).

**Figure 6.**
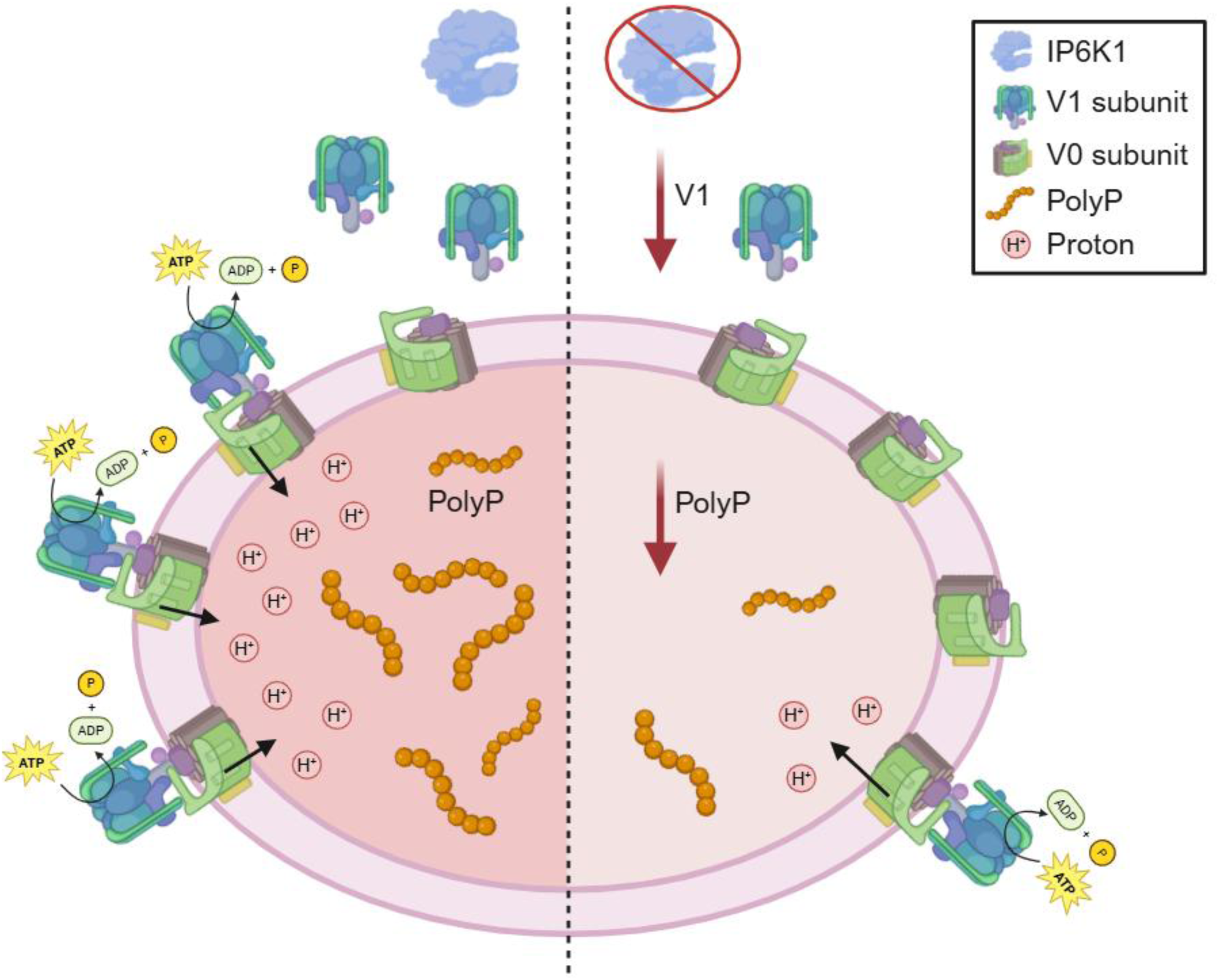
PolyP synthesis in mast cell granules is supported by IP6K1. In RBL-2H3 secretory granules, polyP synthesis is dependent on granule acidification, which is generated and maintained by the activity of V-ATPase. The cytoplasmic catalytic V1 subcomplex of V-ATPase, that hydrolyses ATP to pump protons into the granule, is essential for V-ATPase activity. The loss of IP6K1 selectively reduces the expression of cytoplasmic V1 subunits, thereby lowering V-ATPase activity and resulting in granule deacidification, in turn reducing polyP levels in RBL-2H3 granules.

The absence of a defined polyP synthase has limited our understanding of polyP synthesis and regulation in metazoans. In isolated mammalian mitochondria, polyP production requires an intact potential gradient across the inner mitochondrial membrane, with FoF_1_ ATPase supporting, or directly functioning as, the polyP synthase [45, 47]. In budding yeast, polyP production by the VTC complex and its coupled translocation into the vacuolar lumen, relies on the vacuolar electrochemical gradient [57]. Alkalinization of isolated RBL-2H3 granules by treatment with NH_4_Cl has been shown to negatively affect their polyP retention [15]. On similar lines, our data show that treatment with FCCP to disrupt the proton gradient, or with BafA1 to block V-ATPase activity, abolishes polyP synthesis in isolated mast cell granules. Together, these findings indicate that an intact electrochemical gradient is prerequisite for polyP synthesis in different membrane bound organelles, despite lack of conservation of the enzymes involved in polyP synthesis.

The connection between inositol pyrophosphates and polyP was first uncovered in an *S. cerevisiae* genetic screen, where was seen that deletion of *KCS1*, which encodes the yeast IP6 kinase, nearly abrogated the production of polyP [75]. Subsequent studies revealed that 5-InsP_7_ binding to the SPX domain of VTC4, the catalytic subunit of the VTC complex, allosterically alters VTC4 to facilitate coupled synthesis and translocation of polyP into the yeast vacuole [39, 41–43]. We have previously shown that depletion of 5-InsP_7_ in mammals leads to a reduction in mitochondrial polyP levels and membrane potential [45]. Additionally, IP6K1 was seen to act independently of its catalytic activity to support mitochondrial respiration, which in turn is essential for polyP production in the mitochondria. Mice deficient for IP6K1 also show decreased platelet polyP levels, with associated impairment in blood clotting and a reduction in infection-induced polyP-dependent neutrophil platelet aggregates [44, 62]. In the present study, we observe a similar loss of polyP in an IP6K1 depleted mast cell line and in BMMCs from *Ip6k1^-/-^* mice. In contrast to the regulation of mitochondrial membrane potential and polyP levels which require 5-InsP_7_ synthesis by IP6K1, polyP accumulation and maintenance of the electrochemical gradient in mast cell granules are supported by IP6K1 independent of its catalytic activity. By modulating the expression of V_1_ subunits of V-ATPase, and consequently controlling granule acidification, IP6K1 is able to not only regulate granule polyP levels but also the accumulation of other mediators, including serotonin and tryptase, positioning this InsP kinase as a master regulator of mast cell proinflammatory action. As granule acidification is primarily regulated by the reversible association of the cytoplasmic V_1_ subcomplex with the membrane anchored V_0_ subcomplex to form a functional V-ATPase [76], by regulating the availability of V_1_ subunits IP6K1 may act in both platelet dense granules and mast cell granules to regulate polyP synthesis and accumulation. Together, these data highlight IP6K1 as a regulator of polyP across organelles and organisms.

Despite the discovery of polyP enriched granules in mammalian platelets more than two decades ago [13], the identity of the polyP synthase in these LROs remains elusive. Given the distant shared lineage of megakaryocytes and mast cells, and the common acidocalcisome-like properties of platelet dense granules and mast cell granules, the mechanism of polyP synthesis is likely to be conserved in LROs in both cell types. In contrast to granule polyP synthesis, which requires ATP as a substrate, we have shown that mitochondria use Pi as a substrate for polyP synthesis [45], raising the possibility that different synthases may operate in mitochondria and granules to produce organelle-specific polyP. Whereas mitochondrial FoF_1_ ATP synthase, with its F_1_ ‘head’ oriented towards the mitochondrial matrix, is topologically positioned to support intra-mitochondrial polyP synthesis, V_0_V_1_ ATPase, with the V_1_ subcomplex in the cytoplasm, is precluded from directly supporting polyP synthesis in the granule lumen (Fig. 6). Instead, the essentiality of V_0_V_1_ ATPase in the synthesis of granule polyP is more likely based on its role in sustaining the proton gradient across the granule membrane. How this potential difference supports granule polyP synthesis is unclear – it may provide the necessary energy for ATP-dependent polyP synthesis by an intra-granule polyP synthase, facilitate polyP transport into the granule if synthesis is on the granule membrane, or support polyP retention within the organelle. Future research is likely to provide clarity on the location, machinery, and regulation of granule polyP synthesis in mammals.

## MATERIALS AND METHODS

### Reagents

Unless specified, all chemicals used in this study were procured from Sigma-Aldrich. The primary antibodies used for immunofluorescence (IF), and immunoblotting (IB) along with antibody dilution for each application, and supplier (including catalogue number) are as follows: anti-IP6K1 (Santa Cruz E-11, sc-376290; IB 1:1500), anti-GAPDH (Cloud Clone, CAB932Hu22; IB 1:8000), anti-LaminB1 (Abcam, Ab16048; IB 1:10,000), anti-ATP5A (Abcam, Ab14748; IB 1:10,000), anti-GM130 (BD Biosciences, 610823; IB 1:1000), anti-Calreticulin (CALR; BD Biosciences, 612136; IB 1:1000), anti-Serotonin (Abcam, Ab66047; IF 1:500), anti-V5 tag (Abcam, Ab53418, IB 1:5000), anti-ATP6V0A (Novus Biological, NB1-89342; IB 1:2000), anti-ATP6V1A (Novus biological, NBP1-33021; IB 1:500), anti-ATP6V1B (ABclonal, A6876; IB 1:1000), anti-ATP6V1D (ABclonal, A12940; IB 1:1000), anti-ATP6V1E (ABclonal, A3756; IB 1:1000), anti-Tryptase (MCP-7; Cell Signalling Technology, E9R8A; IB 1:1000; IF 1:500), anti-HEXA (ABclonal, A5646; IB 1:1000; IF 1:250), anti-rabbit IgG(H+L), Alexa Fluor 488 goat (Invitrogen, A11034), anti-rabbit IgG(H+L), Alexa Fluor 555 goat (Invitrogen, A21429), anti-goat IgG(H+L), Alexa Fluor 488 dockey (Invitrogen, A11031), anti-mouse IgG(H+L), Alexa Fluor 488 goat (Invitrogen, A11017), anti-mouse IgG(H+L), Alexa Fluor 568 goat (Invitrogen, A11034). Restriction enzymes were purchased from New England Biolabs. LysoSensor Green DND-189 (L7535), Pierce™ BCA Protein Assay Kit (1856210), Dulbecco’s Modified Eagle’s Medium (DMEM), Fetal Bovine Serum (FBS) and other cell culture reagents were from Thermo Fisher Scientific. PVDF membrane and ECL prime chemiluminescence substrate were procured from GE Healthcare. Protease inhibitor cocktail (PIC; Sigma P8340), and phosphatase inhibitor cocktail 3 (PhosI; Sigma P0044) were procured from Sigma-Aldrich. Plasmids were purified using a plasmid isolation kit (MN NucleoSpin, 740588).

### Cell lines

The RBL-2H3 cell line was procured from ATCC, and the HEK293T cell line, previously authenticated in [45], was provided by Dr Solomon H Snyder (Johns Hopkins School of Medicine, USA). RBL-2H3 cells were grown in a humidified incubator with 5% CO_2_ at 37°C in Dulbecco’s modified Eagle’s medium (DMEM) supplemented with 15% fetal bovine serum (heat-inactivated at 56°C for 30 min; FBS-HI; Thermo Fisher Scientific, 16000044), 1 mM L-glutamine, 100 U/mL penicillin, and 100 µg/mL streptomycin.

The RBL-2H3 non-targeted control cell line and IP6K1 knockdown cell lines were generated using the protocol provided by Addgene (https://www.addgene.org/protocols/plko/). Plasmids encoding the shRNA constructs were generated in the lentiviral vector pLKO.1 (Addgene, 8453) backbone, for a non-targeting control shRNA (Addgene, 1864) or two different shRNAs directed against the 3’UTR of rat IP6K1 (5′-GGACCATTGCTCTGAACTT-3′ and 5′-GCTCTGAGAAGATCAGCT-3′). HEK293T cells were transfected with pLKO.1 encoding the shRNA, along with VSV-G (Addgene, 138479) and psPAX2 (Addgene, 12260), using polyethyleneimine (PEI; Polysciences, 23966) in a ratio of 1:3 (DNA: PEI), and incubated at 37°C to generate lentivirus particles. After 48 h, the culture supernatant was collected and filtered through a 0.45 μm pore-size syringe filter (Whatman, 99132504) to harvest viral particles. The viral suspension was supplemented with polybrene (8 μg/mL, Sigma-Aldrich) and added to RBL-2H3 cells. After 16 h, the media was replaced, and cells were incubated further for 48 h. The transduced cells were selected for puromycin resistance by culturing them for five days in 15% FBS-HI containing DMEM supplemented with puromycin dihydrochloride (2 μg/mL; Sigma-Aldrich, P8833). To verify the extent of IP6K1 knockdown, RBL-2H3 cells were lysed in cell lysis buffer (50 mM HEPES pH 7.4, 100 mM NaCl, 1 mM EDTA, 0.5% Nonidet P40, PIC and PhosI) for 1 h at 4°C. The lysate was sonicated at 20% amplitude for 10 secs (Sonics Vibra-cell), and centrifuged at 18000*g* for 10 min at 4°C. The supernatant was collected and subjected to SDS-PAGE followed by western blot analysis with an IP6K1-specific primary antibody.

Constructs encoding C-terminally V5-tagged catalytically active or inactive (K226A/S334A) mouse IP6K1 [74] for lentivirus-based expression were generated in the plasmid pLX304 (Addgene #25890). Lentivirus particles were prepared in HEK293T cells as described above and used to transduce sh*Ip6k1*-1 RBL-2H3 cells. The transduced cells were cultured for five days in 15% FBS-HI containing DMEM supplemented with blasticidin S HCl (10 µg/mL; Thermo Fisher Scientific, A1113903). A fraction of blasticidin resistant cells was used for western blot analysis to verify the stable expression of IP6K1, and the remaining cells were pooled for cryopreservation.

### Animal experiments

All animal experiments were approved by the Institutional Animal Ethics Committee (Protocol number EAF/RB/01/2025), and were performed in compliance with guidelines provided by the Committee for the Control and Supervision of Experiments on Animals, Government of India (2035/GO/RRcBi/S/18/CCSEA). *Ip6k1^+/+^* and *Ip6k1^-/-^* mice (*Mus musculus*, strain C57BL/6) used for this study were housed in the experimental animal facility at the BRIC-Centre for DNA Fingerprinting and Diagnostics (CDFD), Hyderabad. *Ip6k1^+/+^* and *Ip6k1^-/-^*littermates were produced by breeding *Ip6k1^+/-^* mice and maintained as previously described [77, 78].

### Generation of bone marrow-derived mast cells (BMMCs)

Mouse hematopoietic stem cells were isolated from *Ip6k1^+/+^*and *Ip6k1^-/-^* as described earlier [79]. Briefly, mice were euthanised using CO_2_, and the femurs and tibiae were collected in 10% FBS-containing RPMI media. The muscles surrounding the femurs and tibiae were carefully removed, and the epiphyses were cut from both ends. Bone marrow inside the femur and tibia was expelled by flushing the bones with RPMI using a 10 mL syringe fitted with a 25G needle. The resulting bone marrow suspension was gently crushed and pipetted to break the clumps. The suspension was filtered through a 70 µm cell strainer to dissolve any remaining clumps. The collected cells were centrifuged at 500*g* for 10 min at 25°C. To remove RBC contamination, 1X RBC lysis buffer (155 mM NH_4_Cl, 10 mM KHCO_3_ and 0.1 mM EDTA, pH 7.2) was added and incubated for 2 min at 25°C. 10% FBS-HI containing RPMI was added, and centrifuged at 500*g* for 10 min at 25°C to collect the cells. Approximately 5 x 10^6^ cells were seeded into a T25 flask and cultured for 21 days in 10% FBS-HI containing RPMI supplemented with interleukin-3 (IL-3; 10 ng/ml; PeproTech; 21313). To confirm the differentiation of stem cells to BMMCs, a fraction of cultured cells was collected and examined for expression of the IgE receptor FcεRI, using a flow cytometry-compatible antibody (BioLegend; 134315).

### PolyP extraction and quantification from mammalian cells

PolyP from mammalian cells was extracted using the phenol-chloroform method [45]. The cells were resuspended in TEEL buffer (10 mM Tris-Cl pH 8.0, 10 mM EDTA, 2 mM EGTA, 100 mM LiCl, and 5 mM NaF) and boiled at 95°C for 2 min, followed by sonication at 20% amplitude for 10 sec. Protein in the extract was quantified using the Pierce™ BCA protein assay kit. An equal volume of acid phenol (pH 4.8) was added to the samples and centrifuged at 18,000*g* for 10 min at 25°C. The resulting upper aqueous phase was mixed with an equal volume of chloroform, followed by centrifugation at 18,000*g* for 15 min at 25°C. The top aqueous layer was collected, mixed with 2.5 volumes of absolute ethanol, and incubated at -80°C overnight. Samples were centrifuged at 18,000*g* for 30 min at 4°C. The resulting white or translucent pellet was resuspended in recording buffer (150 mM KCl and 20 mM HEPES, pH 7.0), and polyP levels were quantified using DAPI by the method described in [45]. Briefly, the pellet resuspended in the recording buffer was split into two equal volumes; one was treated with 1 μg of purified recombinant *Saccharomyces cerevisiae* exopolyphosphatase (*Sc*PPX), while the other remained untreated, and both were incubated at 37°C for 16 h. For polyP estimation, 100 μL samples, and polyP_20_ standard (serially diluted, ranging from 312.5 to 10,000 pmoles in Pi equivalents), were transferred to a 96-well black plate (Thermo Fisher Scientific; P8741). An equal volume of 60 µM DAPI (in recording buffer) was added to the samples and standards, and incubated for 20 min at 25°C. DAPI fluorescence was measured using a multimode plate reader (Perkin Elmer EnSpire; Ex 415 nm; Em 550 nm). The amount of polyP in the untreated and *Sc*PPX treated samples were interpolated from a linear regression standard curve of DAPI-PolyP_20_ fluorescence. The difference in the interpolated values corresponded to the amount of polyP in the sample, and was expressed in terms of nmoles Pi/mg protein.

### Immunofluorescence analysis

RBL-2H3 cells were grown on glass coverslips for 24 h. The cells were fixed with 4% paraformaldehyde (PFA) for 10 min at room temperature (RT), and permeabilised with 0.2% Triton-X 100 for 10 min at RT. Non-specific antibody binding was blocked by incubating the cells in blocking buffer (3% BSA in Tris-buffered saline with 0.2% Triton X-100) (TBS-T) for 1 h at RT. For the in situ overlay assay, purified GST-tagged PPBD (5 µg/ml) was diluted in blocking buffer and added to fixed and permeabilised cells for 1 h at RT. The unbound protein was removed by washing thrice with TBS-T for 10 min at RT. Primary antibodies (including anti-GST antibody in case of the in situ overlay) were diluted in blocking buffer and added to the cells for 16 h at 4°C in a humidified chamber. Cells were washed thrice with TBS-T and incubated with fluorophore-conjugated secondary antibodies diluted in blocking buffer for 1 h at RT. Cells were washed with TBS-T, and the coverslips were mounted on glass slides using antifade mounting medium with DAPI (H-1200, Vecta Labs), air-dried and sealed. Images were acquired in a Leica TCS SP-8 confocal microscope using 405, 488, and 514 nm lasers, with a 63x, 1.4 NA objective. All images are z-stacks showing xy dimensions as a maximum intensity projection (MIP). Images were subjected to level adjustment for improved visualisation using LASX (Leica application suite X; Ver-3.4.2.18368). The images were quantified using Fiji software [80].

### LysoSensor Green DND-189 staining

RBL-2H3 cells were seeded in live-cell compatible cell culture dishes for 16 h. Following a rinse with 1X PBS, cells were incubated with LysoSensor Green DND-189 (1 µM; Thermo Fisher Scientific L7535) and DAPI (2 µg/mL) in DMEM for 1 h at 37°C. The cells were washed twice with 1X PBS, and 15% FBS-HI containing DMEM was added to the cells. Live images were acquired in a Leica TCS SP-8 confocal microscope using 405 and 488 nm lasers, and fitted with a 63x, 1.4 NA objective, and presented as described above.

### Subcellular fractionation by differential centrifugation

Approximately 20 x 10^6^ RBL-2H3 cells were harvested in 1X PBS. The cells were pelleted at 300*g* and resuspended in buffer A (50 mM Tris-Cl, pH 7.4, 250 mM sucrose, 5 mM EDTA, 1X PIC, and 1X PhosI), and incubated on ice for 20 min. The cells were lysed by mechanical shearing with 6-8 passes through a 22G needle fitted onto a 5 mL syringe. Lysis was monitored by observing the sample under a phase contrast microscope. The sample was centrifuged at 300*g* for 5 min at 4°C to remove any remaining unlysed cells. The supernatant was centrifuged at 1000*g* for 10 min at 4°C to collect nuclei. The post-nuclear supernatant was centrifuged at 8000*g* for 10 min at 4°C to collect the mitochondrial fraction. The post-mitochondrial supernatant was centrifuged at 12000*g* for 15 min at 4°C to collect the endomembrane system. Finally, the post-endomembrane supernatant was centrifuged at 40,000*g* using a Ti-40 fixed angle rotor in a Beckman Coulter ultracentrifuge to collect the granule fraction. The organelle fractions were suspended in buffer A, and total protein was estimated using the Pierce™ BCA protein assay kit. The samples were subjected to western blotting to probe for organelle-specific protein markers and verify the enrichment of different subcellular fractions.

### Ex vivo polyP synthesis assay

RBL-2H3 cell organelle-enriched fractions were resuspended in reaction buffer (10 mM PIPES-KOH, pH 6.8, 150 mM KCl, and 0.5 mM MgCl_2;_ adapted from [40]), and protein content was quantified using a Pierce™ BCA protein assay kit. The real-time polyP synthesis assay was set up in a 96-well black plate (Thermo Fisher Scientific; P8741) using 20 µg protein along with the indicated concentrations of substrates and inhibitors, and DAPI (30 μM) in a final volume of 100 μL. The reaction mix was incubated at RT for 5 min, and the plate was transferred to a PerkinElmer EnSpire multimode plate reader to record DAPI fluorescence at 1 min intervals for 20 min (Ex 415 nm; Em 550 nm). PolyP synthesis in each sample was quantified as the change in DAPI fluorescence over time, ΔF=F_i_-F_0_ (where F_i_ is the fluorescence at time ‘i’ and F_0_ is the fluorescence at time ‘0’).

### Statistical analysis

GraphPad Prism 8 was used to perform statistical analyses and prepare graphs. Densitometry data for western blots were obtained using Fiji software [80]. Band intensities of indicated proteins were normalised to their respective loading controls from the same blot, and their relative expression was compared in treated or test samples versus untreated or control samples. Individual groups were compared to a control that was normalised to 1 to derive a relative fold-change. All fold change values were transformed to log_2_, subjected to either a one-sample *t-*test against a theoretical mean of 0 or one-way ANOVA with Tukey’s multiple comparisons test, as indicated. For quantification of microscopy images (in arbitrary units), data were analysed by a two-tailed Mann-Whitney test or a Kruskal-Wallis test with Dunn’s multiple comparisons test, as indicated. The number of cells (n) used to perform statistical tests for each experiment, and the number of biologically independent replicates (N) for each experiment, are indicated in the figure legends. All quantified data are presented as mean ± S.D. and individual data points are indicated as scatter dot plots.

## SUPPORTING INFORMATION

Supplementary Figures S1-S4 are provided in a single PDF file.

## DATA AVAILABILITY STATEMENT

All data are available on request.

## AUTHOR CONTRIBUTIONS

**Manisha Mallick**: Conceptualisation, Methodology, Investigation, Validation, Formal analysis, Visualisation, Writing – original draft, Writing – review and editing. **Rashna Bhandari**: Conceptualisation, Writing – review and editing, Supervision, Project administration, Funding acquisition. Authors have read and agreed to the published version of the manuscript.

## ACKNOWLEDGEMENTS

We thank Maddika Subba Reddy, Shweta Tyagi, and members of the Laboratory of Cell Signalling, especially Jayashree Suresh Ladke, for their assistance and valuable feedback; Kuldeep Verma for providing LysoSensor green DND-189; staff at the Sophisticated Equipment Facility and the Experimental Animal Facility at CDFD for technical assistance.

