## Supplementary Figures S1-S4 for "Polyphosphate synthesis in mast cell granules relies on V-ATPase activity supported by IP6K1"

Manisha Mallick: 0009-0004-8971-2510

Rashna Bhandari: 0000-0003-3101-0204

**Running title:** IP6K1 facilitates polyphosphate synthesis in mast cells

**This PDF file includes:**

Supplementary Figures S1-S4

**Figure S1**

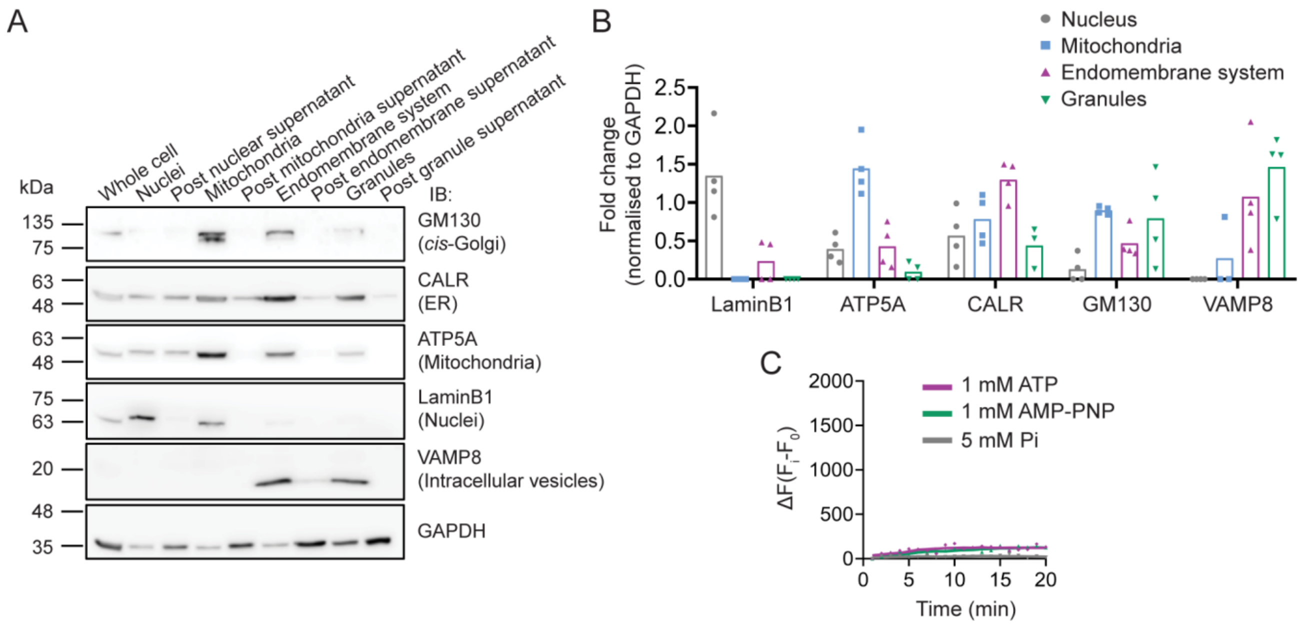


**Figure S1. Subcellular fractionation. (A)** Immunoblots of the indicated cell fractions isolated by differential centrifugation of a lysate from RBL-2H3 cells. The pellets obtained during sequential centrifugation steps yielded organelle enrichment as follows: nuclei (1000*g*), mitochondria (8000*g*), endomembrane system (12000*g*), and granules (40,000*g*). Specific proteins that serve as organelle markers are indicated against each blot. **(B)** Quantification of (A). Scatter bar graphs represent the distribution of Lamin-B1, ATP5A, CALR, GM130 and VAMP8 in nuclei, mitochondria, endomembrane system, and granule fractions isolated from RBL-2H3 cells (N = 4). The level of each indicated protein was normalised to the level of GAPDH for each fraction. The data confirm the enrichment of the nuclear fraction marked by Lamin-B1, mitochondria marked by ATP5A, endomembrane system marked by CALR, and granules marked by VAMP8. **(C)** Control for Fig. 2(A, B). Line graphs represent the change in DAPI fluorescence over time (ΔF) in polyP reaction buffer in the presence of ATP (1 mM), AMP-PNP (1 mM), or Pi (5 mM), confirming that no ΔF is contributed by the substrates alone in the absence of cell fractions.

### Figure S2

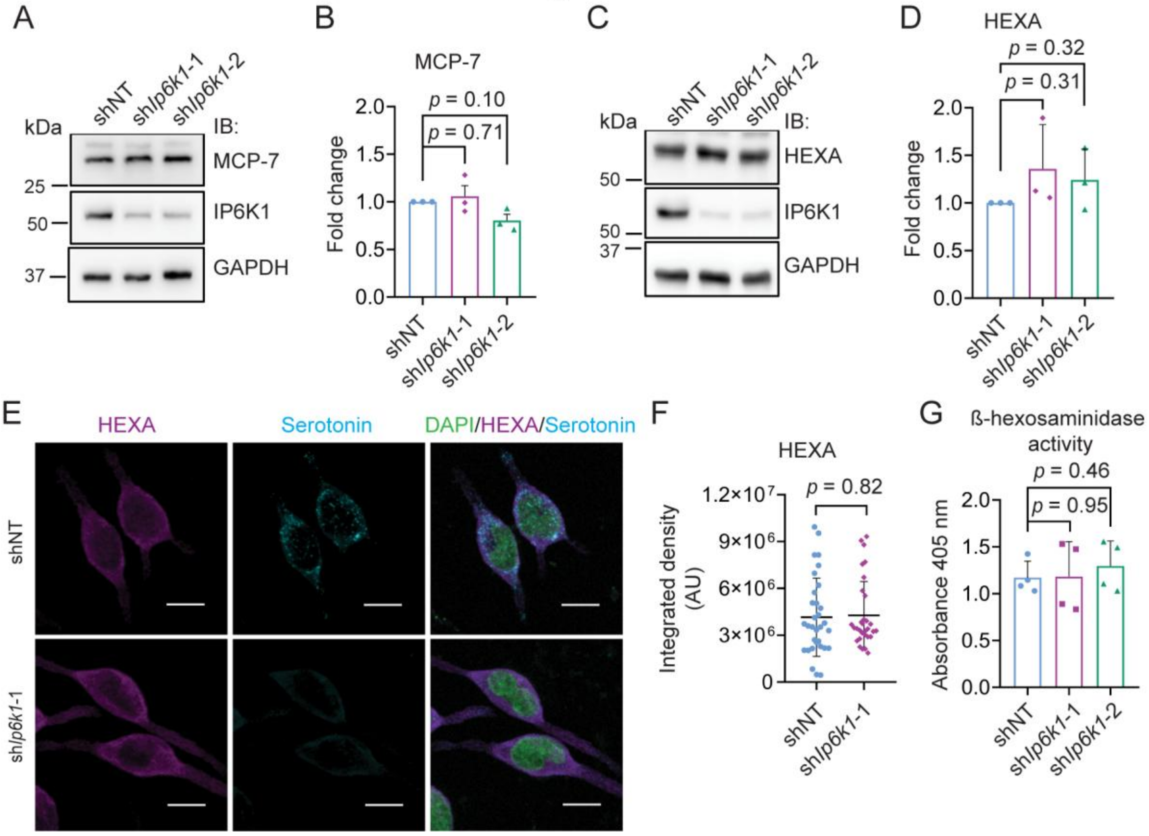


**Figure S2. Effect of IP6K1 depletion on tryptase and β-hexosaminidase. (A)** Immunoblots show the levels of tryptase (MCP-7) and IP6K1 in the indicated RBL-2H3 cell lines, with GAPDH as a loading control. **(B)**. Quantification of (A). Bar graphs (mean ± S.D.) represent levels of tryptase normalised to GAPDH in sh*Ip6k1* cell lines compared with the shNT control cell line (N = 3). The fold change values were transformed to $\log_2$, and *p* values were determined using a one-sample *t*-test against a theoretical mean of 0. **(C)** Immunoblots show the levels of β-hexosaminidase subunit alpha (HEXA) and IP6K1 in the indicated RBL-2H3 cell lines, with GAPDH as a loading control. **(D)**. Quantification of (C). Bar graphs (mean ± S.D.) represent levels of HEXA normalised to GAPDH in sh*Ip6k1* cell lines compared with the shNT control cell line (N = 3). *p* values were determined as in (B). **(E)** Immunofluorescence analysis in the indicated RBL-2H3 cell lines show HEXA (magenta), serotonin (cyan), and DAPI to mark nuclei (green). Images showing individual channels and an overlay were captured with a Leica TCS SP8 confocal microscope using a 63x, 1.4 NA oil-immersion objective, and are presented as z-stacks with xy dimensions as a maximum intensity projection (MIP); scale bar: 10 μm. **(F)** Quantification of (E). Scatter dot plots (with mean ± S.D.) represent HEXA fluorescence signals as integrated density (in arbitrary units, AU) per cell in shNT (n = 32) or sh*Ip6k1*-1 (n = 31) RBL-2H3 cell lines. Data were compiled over three independent experiments. *p* value was determined using a Mann-Whitney test. **(G)** Bar graphs (mean ± S.D.) represent β-hexosaminidase activity in the indicated RBL-2H3 cell lines (N = 4). *p* values were determined using a two-tailed unpaired Student's *t*-test.

#### Figure S3

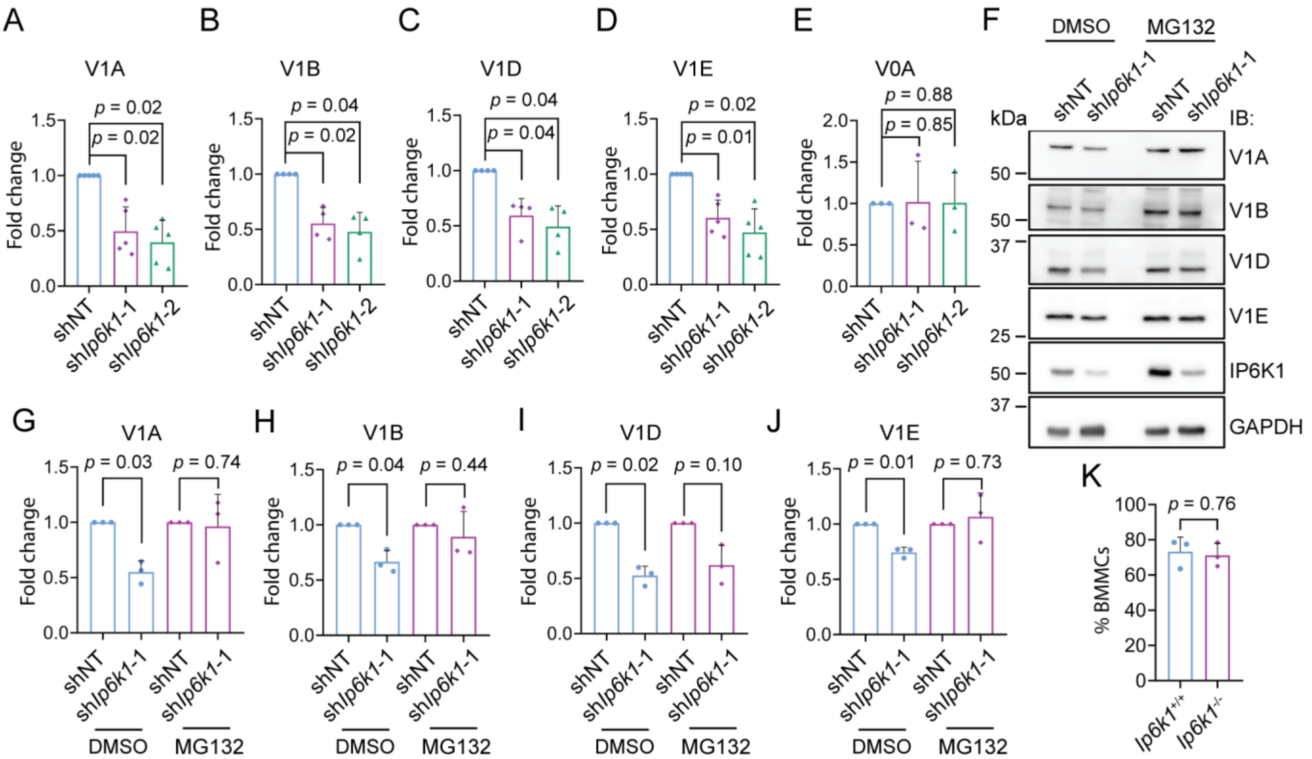


**Figure S3. Loss of IP6K1 downregulates the levels of V1 subunits of V-ATPase. (A-E)** Quantification of Fig. 4(C-E). Bar graphs (mean ± S.D.) represent the levels of V1A (A), V1B (B), V1D (C), V1E (D) and V0A (E) subunits of V-ATPase in sh*Ip6k1*-1 and sh*Ip6k1*-2 RBL-2H3 cell lines, compared with the control (shNT). (N = 5 for A-D; N = 3 for E). The fold change values were transformed to $\log_2$, and *p* values were determined using a one-sample *t*-test against a theoretical mean of 0. **(F)** Immunoblots show the levels of V1A, V1B, V1D, and V1E subunits of V-ATPase in the indicated RBL-2H3 cell lines treated with DMSO or MG132 (10 µM). Representative immunoblots are compiled from different individual blots, each run with a loading control. Representative blots are presented for IP6K1 and the loading control (GAPDH). **(G-J)** Quantification of (F). Bar graphs (mean ± S.D.) show the levels of V1A (G), V1B (H), V1D (I), and V1E (J) subunits of V-ATPase in sh*Ip6k1*-1 RBL-2H3 cells compared with shNT (control), in the presence of DMSO or MG132 (10 µM) (N=3). *p* values were determined as in (A-E). **(K)** Bar graphs (mean ± S.D.) represent the percentage of bone marrow derived mast cells (BMMCs) obtained post-differentiation of bone marrow stem cells isolated from *Ip6k1*$^{+/+}$ and *Ip6k1*$^{-/-}$ mice (N = 3). *p* value was determined using a two-tailed unpaired Student's *t*-test.

**Figure S4**

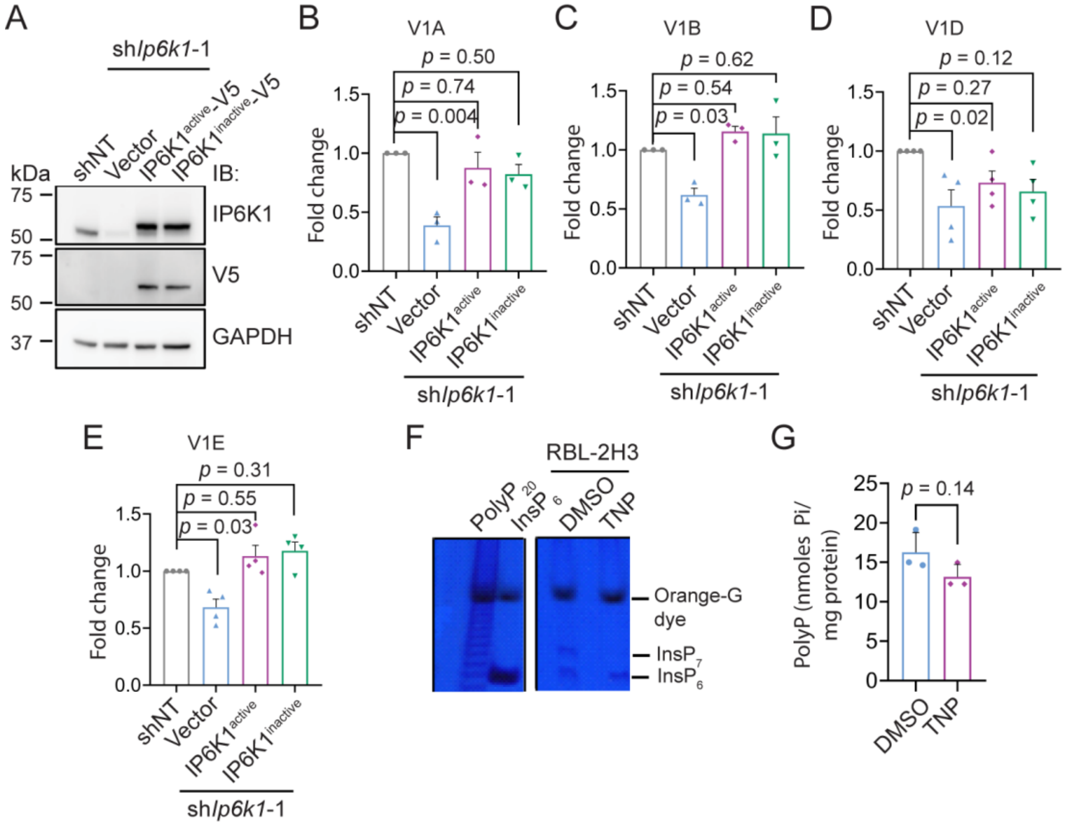


**Figure S4**. **Overexpression of IP6K1 restores V-ATPase V1 subunit levels. (A)** Immunoblots show the levels IP6K1 in non-targeted control (shNT), sh*Ip6k1*-1 transfected with vector alone, or sh*Ip6k1*-1 RBL-2H3 cells stably expressing V5-tagged IP6K1 that is active (IP6K1$^{active}$) or catalytically inactive (IP6K1$^{inactive}$; K226A/S334A). The blots also detect the V5 tag on overexpressed IP6K1, and GAPDH as a loading control. **(B-E)** Quantification of Fig. 5(A, B). Bar graphs (mean ± S.D.) represent the levels of V1A (B), V1B (C), V1D (D) and V1E (E) subunits of V-ATPase in the indicated RBL-2H3 cell lines, compared with shNT (control) (N = 3 for V1A and V1B; N = 4 for VID and V1E). The fold change values were transformed to $\log_2$, and *p* values were determined using a one-sample *t*-test against a theoretical mean of 0. **(F)** Detection of inositol polyphosphates ($InsP_6$ and $InsP_7$) from DMSO or TNP (10µM for 16 h) treated RBL-2H3 cells, enriched using titanium dioxide ($TiO_2$) beads, resolved by 35.8% PAGE and stained with toluidine blue as described in [Wilson, M.S.C. and Saiardi, A. (2018) Inositol phosphates purification using titanium dioxide beads. *Bio Protoc*. 8, e2959]. $PolyP_{20}$ was used as a marker, synthetic $InsP_6$ was used as a standard, and Orange G was used in the loading dye. Non-essential lanes were removed from the same gel for representation. **(G)** Bar graphs (mean ± S.D.) represent the levels of polyP extracted from RBL-2H3 cells treated with DMSO or TNP (10 µM for 16 h). *p* value was determined using a two-tailed unpaired Student's *t*-test (N = 3).
